# OpenSIM: open source microscope add-on for structured illumination microscopy

**DOI:** 10.1101/2023.06.16.545316

**Authors:** Mélanie T.M Hannebelle, Esther Raeth, Samuel M. Leitao, Tomáš Lukeš, Jakub Pospíšil, Chiara Toniolo, Olivier F. Venzin, Antonius Chrisnandy, Prabhu Prasad Swain, Nathan Ronceray, Matthias P. Lütolf, Andrew C. Oates, Guy M. Hagen, Theo Lasser, Aleksandra Radenovic, John D. McKinney, Georg E. Fantner

## Abstract

Super-resolution techniques expand the abilities of researchers who have the knowledge and resources to either build or purchase a system. This excludes the part of the research community without these capabilities. Here we introduce the openSIM add-on to upgrade existing optical microscopes to Structured Illumination super-resolution Microscopes (SIM). The openSIM is an open-hardware system, designed and documented to be easily duplicated by other laboratories, making super-resolution modality accessible to facilitate innovative research. The add-on approach gives a performance improvement for pre-existing lab equipment without the need to build a completely new system.

## Main

The complexity and cost of a scientific instrument have a large impact on the number of scientific teams with access to the instrument for their research^1,2^ and thus to which extent a microscope technique will contribute to making scientific discoveries.

Super-resolution microscopy has become a powerful tool enabling researchers to study biological processes with an exceptional resolution by overcoming the diffraction limit of light inherent in traditional imaging methods. Structured illumination microscopy^3^ (SIM) is a super-resolution technique particularly suited for biological applications due to reduced phototoxicity and high-resolution^4–6^. SIM increases microscopy resolution by using a spatially structured pattern to illuminate the sample, in combination with image reconstruction to deduce information that would normally be unresolved. Although it provides a moderate increase in resolution compared to other super-resolution techniques such as STED^7^ or SMLM^8,9^ (two-fold maximum increase in lateral resolution for linear SIM^3^), biological imaging benefits from higher viability when utilizing SIM as it requires lower light excitation intensity resulting in reduced phototoxicity and photobleaching compared to other techniques^1^.

The hurdle for most research groups is the limited access to super-resolution microscopes, which forces them to rely on lower-resolution conventional fluorescence microscopes instead. To promote technology dissemination among researchers, open-source microscopy initiatives have an significant impact on the distribution of optical designs and instrument building techniques^10,11^. Several initiatives have introduced stand-alone research-grade open-source microscopes, among them the light-sheet microscope openSPIM^10,12^, the wide-field microscope OpenFlexure^13^, the 3D-printed optical toolbox UC2^14^, the AttoBright^2^ for single-molecule detection, and the miCube^15^ for single molecule localization microscopy. Although open-source microscopes have provided scientists with an easier access to super-resolution imaging methods, there is a lack of open-source integrative systems which allow users to extend commercial microscopes readily available in most labs.

Here we introduce the openSIM, a microscope add-on that extends the imaging capabilities of already existing microscopes without the need to build a completely new system. At the same time, this add-on does not impact the function of additional instrumentation connected to the microscope, such as incubation chambers to control temperature, CO2 and humidity, microscope stages, syringe pumps, microfluidic systems etc. depending on the samples being studied. We believe that this reduces the entry barrier to adopt open source super-resolution technology.

As an open source microscope, we provide detailed documentation and instructions^16^ enabling other scientists to build their own openSIM add-on (Supplementary Figure 1, Supplementary Figure 2) to upgrade their microscope setups into super-resolution instruments, and benefit from improved image datasets for their biological research. The assembled openSIM add-on is a compact module (Figure 1a) that connects to the illumination port of standard fluorescence microscopes (Figure 1b, Supplementary Figure 3). It replaces the illumination source of the microscope and provides the striped illumination pattern required for SIM imaging (Supplementary Figure 4). For this, collimated light from high-power LEDs illuminates the spatial light modulator which generates the pattern to be projected on the sample via the openSIM tube lens and the microscope objective (Figure 1c, Figure 1d). A software interface brings together on-the-fly pattern control, illumination color control, light intensity control, camera control (exposure time, camera gains, data saving etc.), and closed loop control of the openSIM heat sink temperature (Supplementary Figure 5). An interface box containing a DAQ (Data Acquisition) device and connectors allows simple cabling between the elements of the system (openSIM add-on, camera, computer and optionally microscope z-stage for automated volumetric imaging) (Figure 1a, Supplementary Figure 6). The openSIM software guides the user to tune the optimal pattern size, depending on the chosen microscope objective. The images acquired with patterned illumination are saved in a format directly compatible with SIMToolbox^17^, an open source MATLAB based software for SIM image reconstruction.

**Figure 1:**
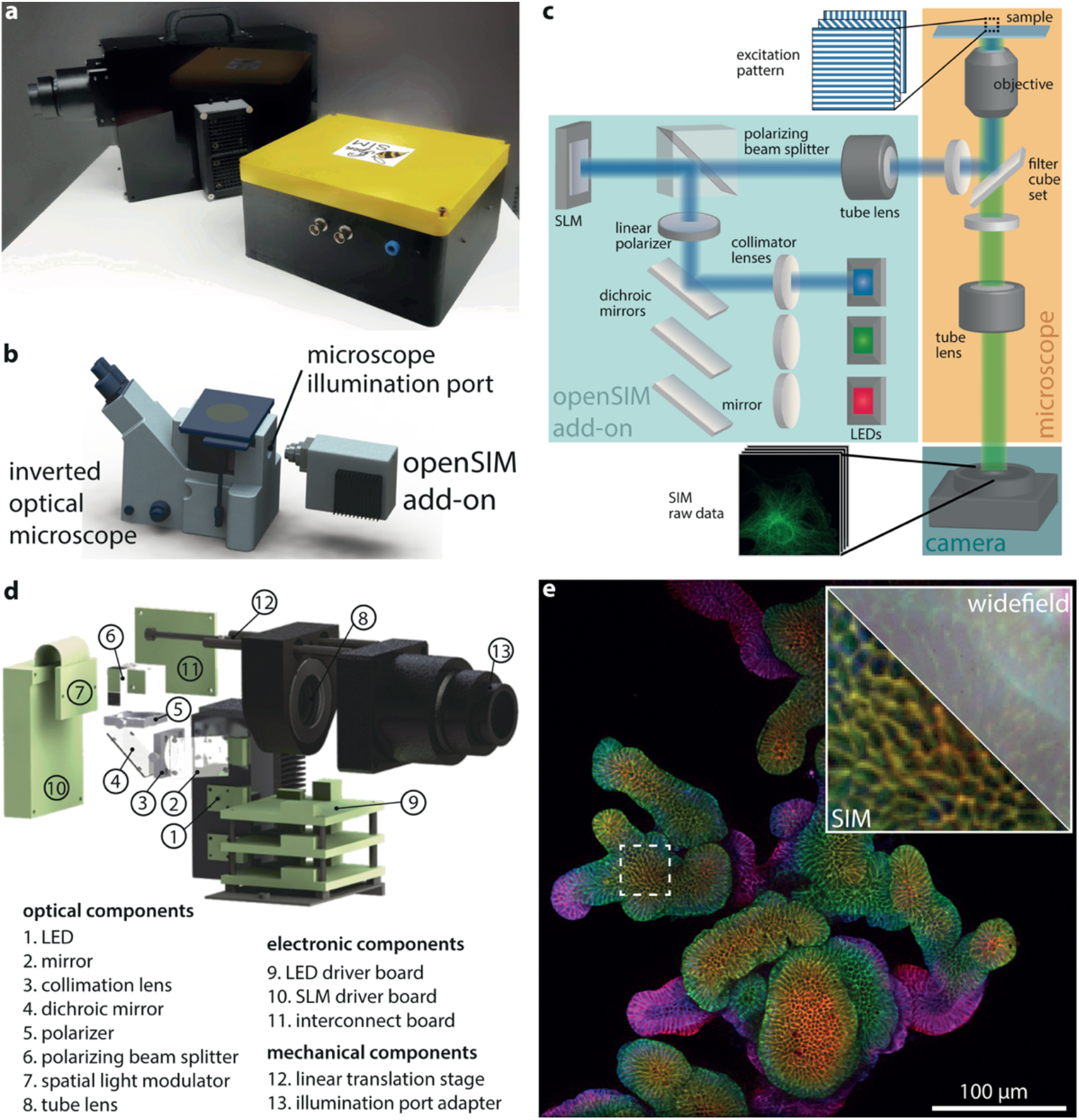
Design and assembly of the openSIM microscope add-on. (a) Photo of the openSIM add-on (back) and its interface box (front). (b) 3D rendering illustrating the connection between the openSIM and a standard inverted optical microscope. (c) Schematic of the optical design of the openSIM add-on. LEDs are collimated to illuminate the spatial light modulator (SLM). The SLM generates a pattern by changing the polarity of the reflected light. The illumination pattern is transferred into the infinity space with a tube lens. (d) 3D rendering of the main optical, electrical and mechanical components of the openSIM. For simplicity only the components associated with the blue illumination channel are represented. (e) Illustration of the increased resolution and optical sectioning when using openSIM compared to wide-field imaging. Whole mount 3D color coded image of fixed mouse intestinal organoids labeled for E-cadherin (epithelial cell junctions). The inset represents a zoom in the area highlighted with a dotted square. Part of the image is not processed (equivalent wide-field image, top right corner), while the bottom left corner is a SIM reconstruction.

**Figure 2:**
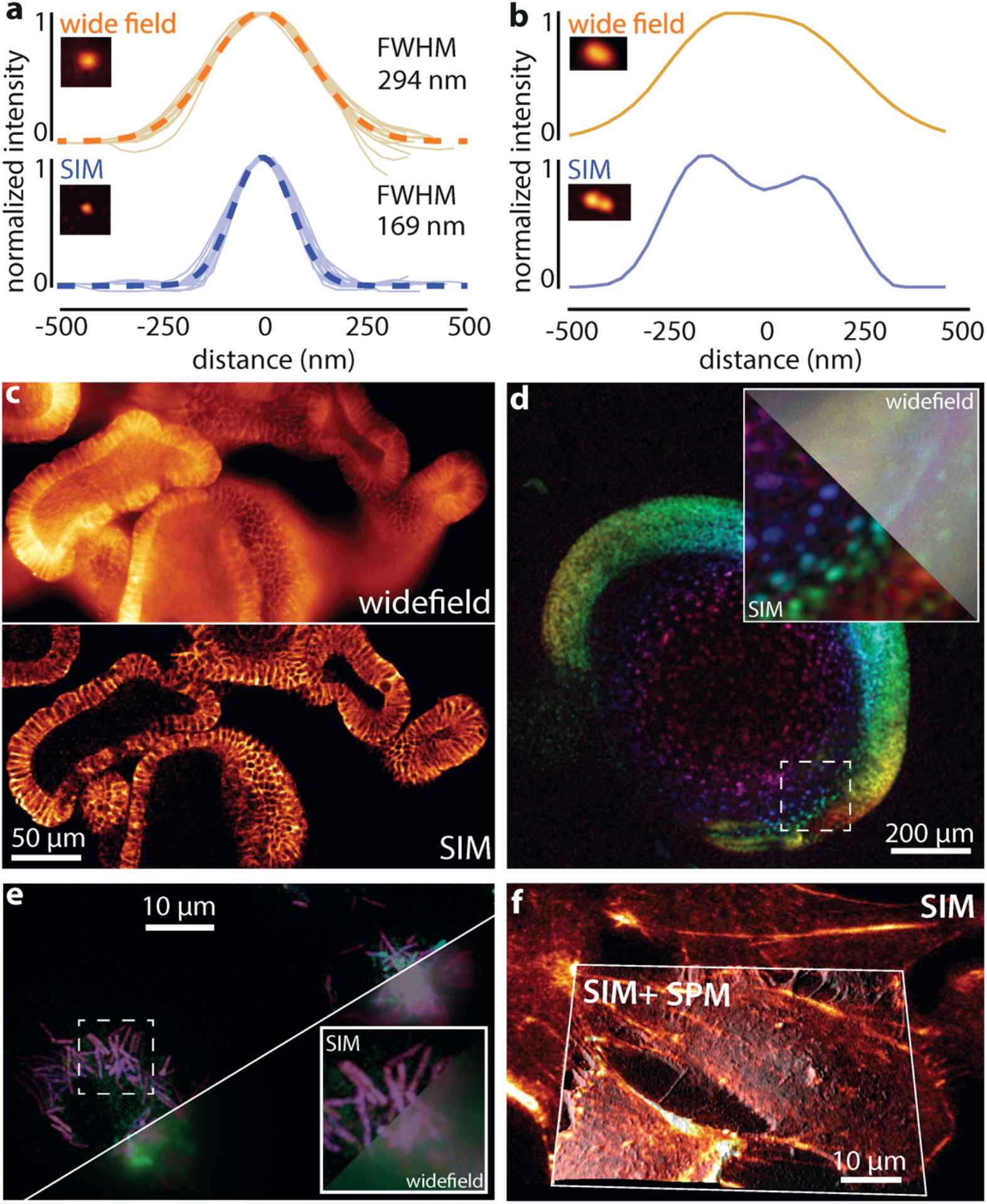
openSIM increases resolution and optical sectioning compared to wide-field illumination. (a) Comparison between the point spread function of 20 isolated 100 nm beads, with wide-field illumination (top) and with the openSIM (bottom), with blue illumination. FWHM: full width at half maximum. Dotted line: average Gaussian fit. The picture inset shows a representative 100nm bead. (b) Comparison between the openSIM and wide-field illumination to resolve two individual beads placed close to each other. (c) Illustration of the optical sectioning effect when using the openSIM. On the wide-field image, out of focus light is noticeable in the image; on the openSIM image, only the part of the sample directly in focus is visible. Sample: fixed mouse intestinal organoids labeled for E-cadherin. (d) 3D color coded image of a live zebrafish embryo with cell nucleus staining. The inset represents a zoom in the area highlighted with a dotted square. Part of the image is not processed (equivalent wide-field image, top right corner), while the bottom left corner is a SIM reconstruction. (e) 3D color-coded image of fixed macrophages infected with *Mycobacterium smegmatis* bacteria. (f) Combined SIM and scanning probe microscopy (SPM) image of fixed actin-stained COS-7 cells. The SIM image is a maximum intensity projection, overlaid as a skin on the topography image obtained with a scanning ion conductance microscope.

To evaluate the performance of the openSIM, we imaged 100 nm fluorescent beads (Supplementary Figure 7, Supplementary Figure 8). After SIM reconstruction, the width of the point spread function was 169 nm (average value on 20 beads), compared to 294 nm for wide-field illumination (Figure 2a). Beads in close proximity that were not resolved with wide field microscopy were resolved with openSIM (Figure 2b): while one single large peak was visible on the wide-field data, two distinct peaks were observed on the SIM image. In addition to increasing the lateral resolution of the microscope, SIM provides an axial optical sectioning effect, more precisely, it reduces the contribution of out-of-focus light to the image. In Figure 2c, we compare a wide-field image and an openSIM image of a fixed mouse intestinal organoid. While out-of-focus cells create a blurred background and lower the contrast in the wide-field image, the openSIM image only contains contributions from cells in the focal plane, thereby greatly increasing the image quality.

To demonstrate the usage of openSIM across a wide spectrum of biological applications, we selected a range of biological samples encompassing large-scale biological specimens as well as samples at cellular and subcellular levels. We imaged a mouse intestinal organoid (Figure 1e), a valuable tool for studying intestinal diseases and interactions between the epithelial tissue, immune system and microbiome. The openSIM add-on also benefits studying whole model organisms, such as zebrafish embryos (Figure 2d). In both samples, the advantages of the optical sectioning are particularly evident. The openSIM was also employed to examine structures at a cellular and subcellular scale, such as the distribution and organization of individual *Mycobacterium smegmatis* bacteria within macrophages during infection (Figure 2e). Tubulin was imaged on a fixed pulmonary artery endothelial cell, using the blue excitation channel of the openSIM (Supplementary Figure 9); this shows the benefit of the openSIM add-on for studying mechanobiology at the subcellular scale. We also mounted the openSIM add-on to a scanning ion-conductance microscope (SICM)^18^, which is a scanning probe microscopy (SPM) technique, for correlated imaging of the cell surface with SICM and the cytoskeleton with the openSIM (Figure 2f). This application further highlights the versatility of the openSIM add-on, showcasing its ability to seamlessly integrate with complex instruments without compromising their performance.

In addition to the two main factors limiting the quality of microscopy images - diffraction of light and limited photon budget - the complexity and cost of an instrument significantly impact the extent to which a microscopy technique will contribute to new scientific discoveries. Our goal in designing the openSIM add-on was not to compete with the most advanced SIM systems that use coherent light from lasers and generate the pattern through interference which yields a higher resolution improvement, due to a higher achievable pattern contrast and overall illumination intensity. Instead, our goal in designing the openSIM add-on was to improve existing instruments without too many modifications and at a moderate cost for components. LEDs sources, in combination with a spatial light modulator allow for a much simpler optical design, therefore the openSIM relies on incoherent LED light sources, and the pattern is generated by projection rather than interference. We leveraged technology initially developed for picoprojectors, as these components are readily available at moderate cost from electronics supply companies and have shown good performance for SIM imaging^16^. This design has the advantage of providing reliable performance, in addition to lower damage due to phototoxicity and bleaching while requiring minimal alignment and maintenance.

The openSIM quantitatively improves the quality of microscopy images (Figure 2), both in terms of resolution (Figure 2a, Figure 2b) and optical sectioning (Figure 2d). Improving the capabilities of existing research grade instruments through the open-hardware add-on approach allows researchers to increase the performance of their instruments for a cost less than one order of magnitude lower compared with commercial SIM microscopes. The openSIM upgrades these dedicated and customized microscopes without reducing their original capabilities, since the setup remains compatible with the specialized equipment often required for studying biological organisms. As the openSIM is a versatile tool with a well-documented assembly, simple maintenance, and linked to an open-source community, we believe that it has the potential to make super-resolution microscopy not only a cutting-edge technique but also a daily tool for biological research.

## Methods

The optical, mechanical and electronic parts of the openSIM are depicted in Supplementary Figure 1. The estimated cost for all components necessary to build up an openSIM add-on is around 10.000 €. A detailed documentation, assembly and user instructions can be found on our wiki-page (https://opensim.notion.site/).

### Optical components for the openSIM

Three high power PT54 LEDs (Luminous Devices) are used to generate the illumination light. The light emitted by the three LEDs is deflected by three mirrors (#43-875, Edmund optics), is collimated using 50 mm lenses (#66-018, Edmund optics), and combined into one optical path with dichroic mirrors (#69-898, #69-900, Edmund optics). A polarizer selects for the specific polarization reflected by the polarizing beam splitter (#49-002, Edmund optics). Then, the light reaches the spatial light modulator (QXGA-3DM-STR, ForthDimension Displays). A tube lens (TTL180-A, Thorlabs), compatible with the commercial microscope being used in this paper (IX71 and IX81, Olympus), is placed in such a way that the SLM is in the back focal place of that tube lens. Commercially available connectors (SM2Y1, LCP11, Thorlabs) were used to connect the openSIM to the illumination port of the microscope. The openSIM is designed to be easily adaptable to other commercially available microscopes (e.g. Olympus, Zeiss, Leica) by changing the tube lens and the illumination port adapter which are available in the Thorlabs catalog.

### Mechanical components for the openSIM

The openSIM was designed with Solidworks software and was printed with a consumer grade 3D printer with PLA as a printing material.

The position of the tube lens is adjusted using a linear translation stage. The translation stage was built with two linear shafts (SSFJ6-100, Misumi), a precision feed screw (XBRF6, Misumi) and two wire springs (WF8-40).

### Electronic components for the openSIM

The LEDs are powered and controlled using the boards and connectors supplied in the DK114N3 development kit (Luminous devices). A custom interconnect board was designed to facilitate the connection between the elements of the openSIM.

The openSIM interface box allows a simple cabling between the elements of the system (openSIM add-on, camera, computer and optionally z-stage). It contains a DAQ (USB-6000, National Instruments), two power supplies (RD-35A, Mean Well; ZWS300BAF-12, TDK) and several connectors to interface the different components. Several analog and digital input and output lines of the DAQ remain available for customization by users.

### LabView interface and openSIM control

We have designed a complete instrument interface with LabView, which brings together on-the-fly pattern control, illumination color control, light intensity control, camera control (exposure time, camera gains, data saving etc.), and closed loop control of heat sink temperature (Supplementary Figure 5). It also saves the detailed acquisition parameters and pre-formats the data to be compatible directly with the open-source SIMToolbox SIM processing algorithm. The openSIM add-on can also provide homogeneous illumination for wide-field imaging, acting as a conventional 3-color light source if SIM is not needed. The modular design of the software provides the possibility to include the control of other commercial cameras.

### Sample preparation

The organoids were derived from Lgr5–eGFP–ires–CreERT2 mouse intestinal crypts and grown in 3D Matrigel culture under expansion conditions (ENRCV)^19^. The preparation of the imaging sample was adapted from Gjorevski *et al*^20^. In short, the organoids were fixed with 4% paraformaldehyde (PFA) in PBS 1X. The fixed samples were permeabilized with 0.2% Triton X-100 in PBS 1X (1 h, room temperature) and were blocked with 0.01% Triton X-100 in PBS 1X containing 10% goat serum (3 h, room temperature). The organoids were then stained using a monoclonal anti-E cadherin antibody (ab 11512; Abcam) followed by secondary antibody Goat-α-Rat Alexa Fluor 568 (A-11077; ThermoFisher Scientific). Extensive washing steps were performed subsequent to each antibody incubation step. The imaging slide was prepared by mounting the stained organoids on glass coverslips with addition of mounting medium Fluoromount-G^®^ (0100-01; SouthernBiotech).

Bovine pulmonary artery endothelial (BPAE) cells with stained F-actin and tubulin were acquired from Thermofisher (FluoCells prepared slide #2). F-Actin is labeled with Texas RED-X phalloidin, and tubulin is labeled with anti-bovine α-tubulin mouse monoclonal 236-10501 conjugated with BODIPY FL goat anti-mouse IgG antibody.

Bone marrow derived macrophages (BMDMs) were differentiated by seeding 10^6^ bone marrow cells from C57BL/6 mice in petri dishes and maintaining them in DMEM supplemented with 10% heat-inactivated fetal bovine serum (HI-FBS), 1% sodium-pyruvate, 1% GlutaMax and 20% L929-cell-conditioned medium as a source of granulocyte/macrophage colony stimulating factor. After 1 week of cultivation at 37°C with 5% CO_2_, the adherent differentiated macrophages were gently detached from the plate using a cell lifter and resuspended in DMEM supplemented with 5% HI-FBS, 1% sodium-pyruvate, 1% GlutaMax and 5% L929-cell-conditioned medium. 10^4^ cells were seeded in a 35 mm cell culture micro-dish with a coverslip bottom (IBIDI) and allowed to adhere for 24 h before infection.

A GFP expressing *Mycobacterium smegmatis* (*Msm*) strain was cultured at 37°C in Middlebrook 7H9 (Difco) supplemented with 10% ABS, 0.5% glycerol, and 0.02% tylox pol. 1 ml of culture at OD_600_ 0.5 was pelleted, concentrated 5 times in the medium of the BMDMs and filtered with a 5 μm filter to obtain a single cell suspension. BMDMs were infected with the single cell suspension with MOI 1:1 and incubated at 37°C with 5% CO_2_. After 4 h the cells were washed extensively with DMEM to eliminate extracellular bacteria and incubated at 37°C with 5% CO_2_ for 48 h to allow the infection to proceed. Before imaging, the cells were stained with CellMask™ Orange (Invitrogen) according to the manufacturer’s protocol and fixed with 4% formaldehyde in PBS for 30 minutes at room temperature.

African green monkey kidney fibroblast-like cells (COS-7), purchased from ATCC were grown in DMEM without phenol red medium (Sigma Aldrich), containing 10% fetal bovine serum. The #1.5 cover glass coverslips were cleaned with piranha solution and coated with fibronectin from bovine plasma (0.5 μM/ml). Then the cells were fixed with 4% PFA in 1xPBS (pH 7.4) for 10 min at room temperature and washed twice for 5 min with PBS (pH 7.4). Prior to staining, the fixed cells were incubated with PBS containing 1% BSA for 30 min to reduce nonspecific background. To visualize F-actin, the phalloidin staining solution (Fluor 488 phalloidin ThermoFisher) was placed on the coverslip for 20 minutes at room temperature.

Zebrafish were maintained according to standard procedures at the EPFL fish facility, which has been accredited by the the Service de la Consommation et des Affaires Vétérinaires of the canton of Vaud – Switzerland (VD-H23) and embryos were staged according to Kimmel et al^21^. Embryos were obtained from outcrosses of Tg (Xla.Eef1a1:H2B-mCherry) to WT fish by natural spawning and raised in fish water (pH 7.8 ± 0.1, conductivity 500 ± 50 μS). For imaging, embryos were de-chorionated and laterally aligned in conical depressions in a pad of 2% low-melting agarose (Sigma) that was cast in a glass-bottom dish (Matek, 35 mm), as in Herrgen et al^22^.

### Combined SIM/SICM microscopy

Scanning probe microscopy was performed with a custom-made scanning ion conductance microscope^18^ (SICM). The sample was actuated in X and Y by a piezo-stage (Piezosystem Jena TRITOR102SG). The capillary was moved in Z by a home-built actuator, operated in hopping mode. The hopping height was 1 μm at 100 Hz rate. The current setpoint used in the hopping actuation was 99% of the normalized current recorded. Images with 256×256 pixels were generated.

### Image processing

The raw patterned images were used to calculate the SIM image using the SIMToolbox^17^ open-source software. Volumetric images were rendered using maximum intensity projections and color-coded projections, using Fiji^23^. Fiji was also used for measuring the point spread function profiles in Figure 2a. The SICM image with SIM overlay (Figure 2f) was rendered using the Fiji plug-in 3D-surface-plot^24^.

## Author contributions

^*^ M.T.M.H. and E.R. contributed equally to this work.

## Contribution statement

M.T.M.H. and G.E.F. conceptualized the idea, M.T.M.H., E.R. designed the instrument, developed the hardware components and the acquisition software, M.T.M.H designed the experiments, M.T.M.H, E.R. created the project documentation, M.T.M.H., E.R., S.M.L, C.T., O.V., A.C., N.R., prepared the samples and acquired the data, M.T.M.H., E.R., S.M.L analyzed the data, T.L., J.P., P.P.S supported the image reconstruction, M.T.M.H., E.R., S.M.L and G.E.F. wrote the manuscript. G.E.F supervised, conceived, and planned the project. All authors reviewed, edited and approved the manuscript

## Acknowledgement

This research was funded by the EPFL Open Science Fund, the Swiss Commission for Technology and Innovation (CTI-18330.1) and the European Research Council (ERC-2017-CoG; InCell.), Swiss National Science Foundation (205321_134786 and 205320_152675), from the Commission for Technology and Innovation under CTI (18330.1 PFNM-NM). The authors would like to thank Adrien Descloux, David Nguyen and Nicolas Bichon for their valuable contributions.

## Supplementary information

Supplementary information is available for this paper. The open hardware documentation for the OpenSIM is available at the following webpage: https://opensim.notion.site/. The webpage is currently private (only accessible through this link) and will be made public after the review of the manuscript.

## References

1. Schermelleh, L. et al. Super-resolution microscopy demystified. Nat Cell Biol 21, 72–84 (2019).

2. Brown, J. W. P. et al. Single-molecule detection on a portable 3D-printed microscope. Nat Commun 10, 5662 (2019).

3. Gustafsson, M. G. L. Surpassing the lateral resolution limit by a factor of two using structured illumination microscopy. J Microsc 198, 82–87 (2000).

4. Coltharp, C. & Xiao, J. Superresolution microscopy for microbiology. Cellular Microbiology 14, 1808–1818 (2012).

5. Shao, L., Kner, P., Rego, E. H. & Gustafsson, M. G. L. Super-resolution 3D microscopy of live whole cells using structured illumination. Nat Methods 8, 1044–1046 (2011).

6. Demmerle, J. et al. Strategic and practical guidelines for successful structured illumination microscopy. Nat Protoc 12, 988–1010 (2017).

7. Hell, S. W. & Wichmann, J. Breaking the diffraction resolution limit by stimulated emission: stimulated-emission-depletion fluorescence microscopy. Opt. Lett. 19, 780 (1994).

8. Betzig, E. et al. Imaging Intracellular Fluorescent Proteins at Nanometer Resolution. Science 313, 1642–1645 (2006).

9. Rust, M. J., Bates, M. & Zhuang, X. Sub-diffraction-limit imaging by stochastic optical reconstruction microscopy (STORM). Nat Methods 3, 793–796 (2006).

10. Marx, V. Microscopy: OpenSPIM 2.0. Nat Methods 13, 979–982 (2016).

11. Girstmair, J. et al. Light-sheet microscopy for everyone? Experience of building an OpenSPIM to study flatworm development. BMC Developmental Biology 16, 22 (2016).

12. Pitrone, P. G. et al. OpenSPIM: an open-access light-sheet microscopy platform. Nat Methods 10, 598–599 (2013).

13. Sharkey, J. P., Foo, D. C. W., Kabla, A., Baumberg, J. J. & Bowman, R. W. A one-piece 3D printed flexure translation stage for open-source microscopy. Review of Scientific Instruments 87, 025104 (2016).

14. Diederich, B. et al. UC2 – A 3D-printed General-Purpose Optical Toolbox for Microscopic Imaging. in Imaging and Applied Optics ITh3B.5 (2019).

15. Martens, K. J. A. et al. Visualisation of dCas9 target search in vivo using an openmicroscopy framework. Nat Commun 10, 3552 (2019).

16. openSIM documentation https://opensim.notion.site/.

17. Křížek, P., Lukeš, T., Ovesný, M., Fliegel, K. & Hagen, G. M. SIMToolbox: a MATLAB toolbox for structured illumination fluorescence microscopy. Bioinformatics 32, 318–320 (2016).

18. Leitao, S. M. et al. Time-Resolved Scanning Ion Conductance Microscopy for Three-Dimensional Tracking of Nanoscale Cell Surface Dynamics. ACS Nano 15, 17613–17622 (2021).

19. Yin, X. et al. Niche-independent high-purity cultures of Lgr5+ intestinal stem cells and their progeny. Nat Methods 11, 106–112 (2014).

20. Gjorevski, N. et al. Designer matrices for intestinal stem cell and organoid culture. Nature 539, 560–564 (2016).

21. Kimmel, C. B., Ballard, W. W., Kimmel, S. R., Ullmann, B. & Schilling, T. F. Stages of embryonic development of the zebrafish. Developmental Dynamics 203, 253–310 (1995).

22. Herrgen, L., Schröter, C., Bajard, L. & Oates, A. C. Multiple embryo time-lapse imaging of zebrafish development. Methods Mol Biol 546, 243–254 (2009).

23. Schindelin, J. et al. Fiji: an open-source platform for biological-image analysis. Nat Methods 9, 676–682 (2012).

24. Barthel, K. U. 3D-Data Representation with ImageJ. (2006).

