## Supplementary Material for "OpenSIM: open source microscope add-on for structured illumination microscopy"

### Supplementary figures

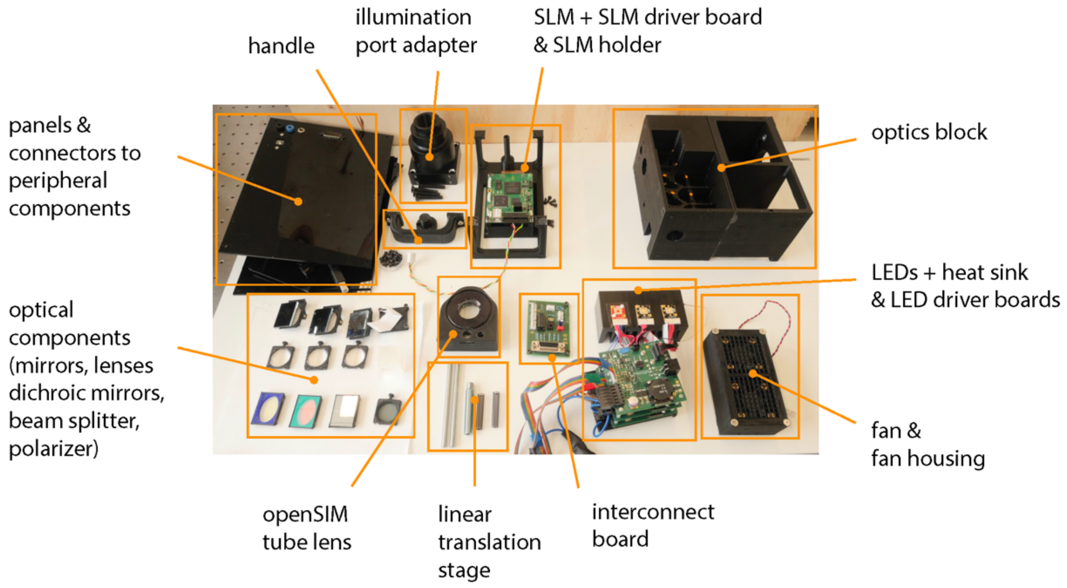


Supplementary Figure 1: Components of the openSIM

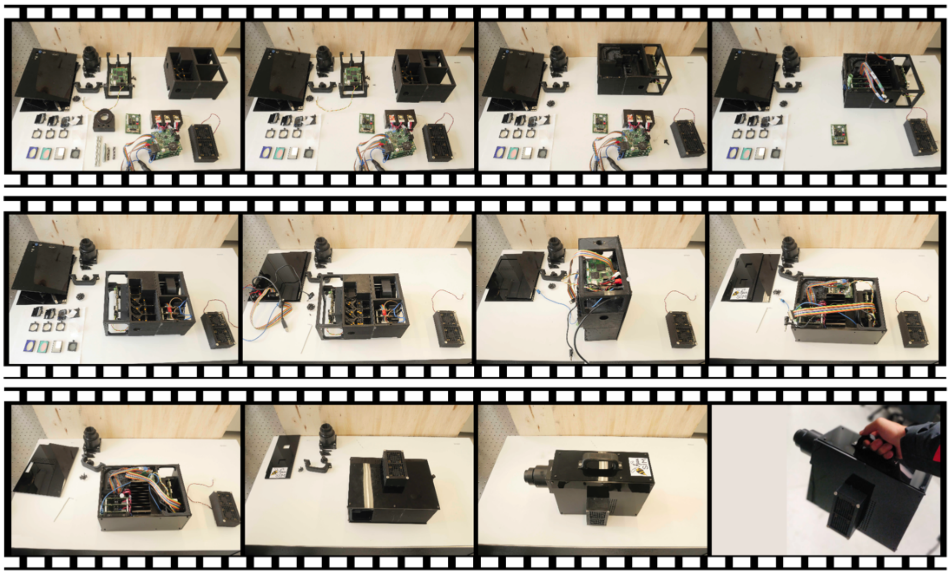

**Supplementary Figure 2: Time-lapse of the assembly of an openSIM**

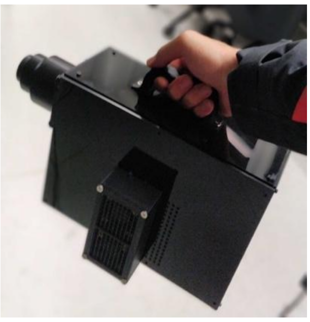

**Supplementary Figure 3: The openSIM is portable and can be moved from one instrument to another**

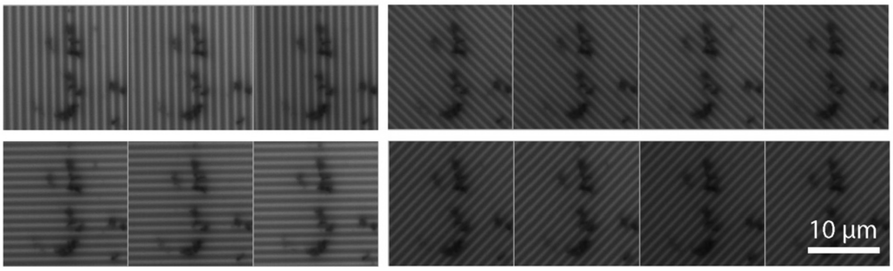


**Supplementary Figure 4: Patterned illumination with the openSIM**

Patterned illumination with the openSIM on a thin fluorescent film. In this example, 4 angles are used and 14 different patterns.

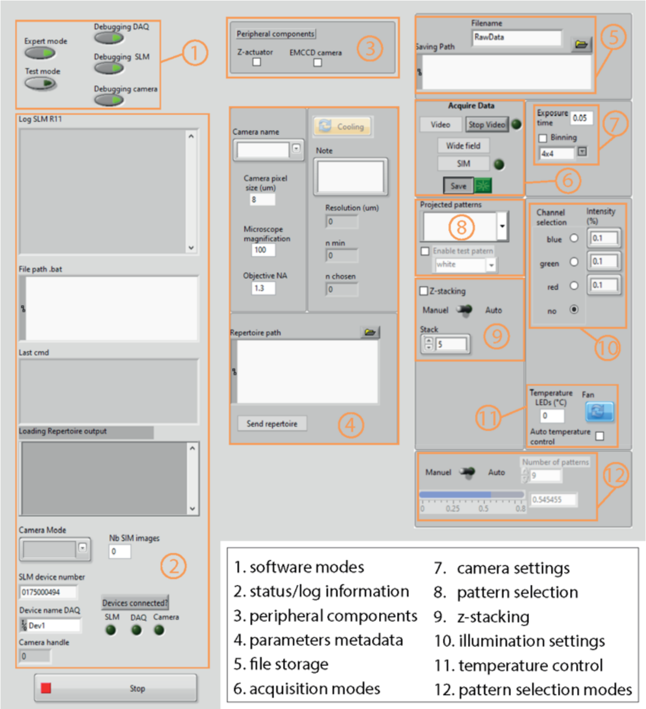


**Supplementary Figure 5: User interface of the openSIM LabView software**

The openSIM software integrates the various components of openSIM including SLM, DAQ and camera, enabling efficient control over pattern generation, illumination management, temperature regulation, and data handling. Its modular architecture enables users with the flexibility to customize and tailor the software to their specific needs and requirements.

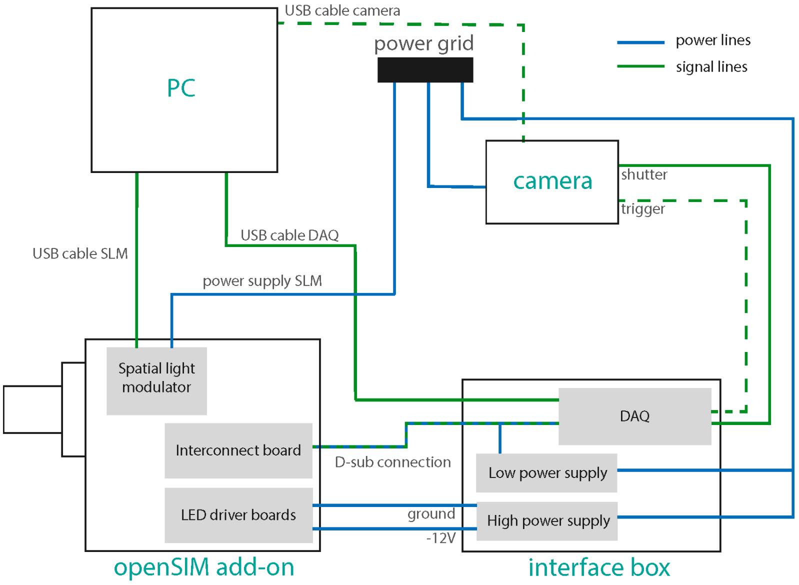


**Supplementary Figure 6: Schematic of the connections between the different components of the openSIM system**

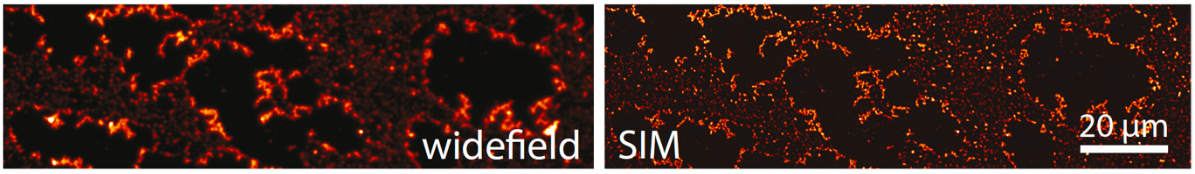


**Supplementary Figure 7: Comparison between SIM and wide-field images of fluorescent beads**

Comparison between an image of 100 nm beads with wide-field blue illumination compared to the openSIM image.

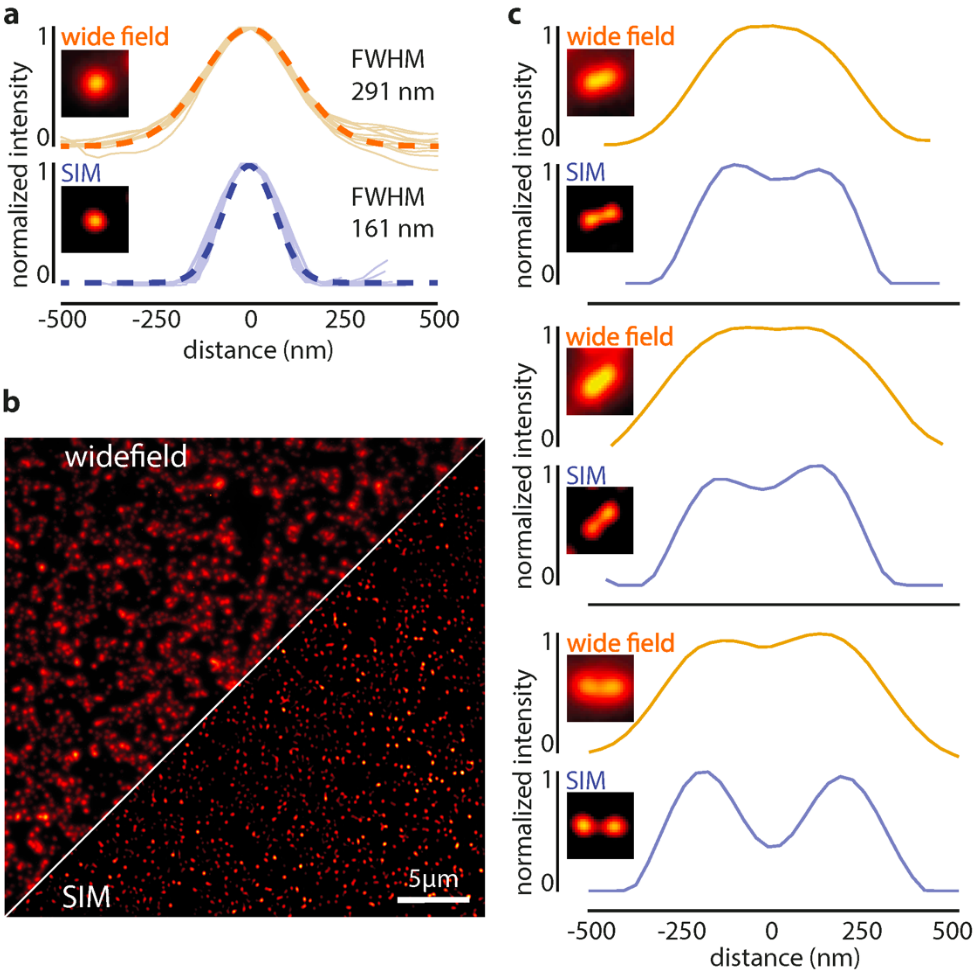


**Supplementary Figure 8: Comparison between SIM and wide-field images of fluorescent beads with openSIM.** Comparison between the point spread function of isolated 100 nm beads (TetraSpeck beads, 100 nm diameter), with wide-field illumination (top) and with the openSIM (bottom), with blue illumination. FWHM: full width at half maximum. Dotted line: average Gaussian fit. The picture inset is a representative point spread function. (b) Comparison between an image of 100 nm beads with wide-field blue illumination compared to the openSIM image. (c) Comparison of the resolving performance between the openSIM and wide-field illumination of two individual 100 nm beads placed close to each other.

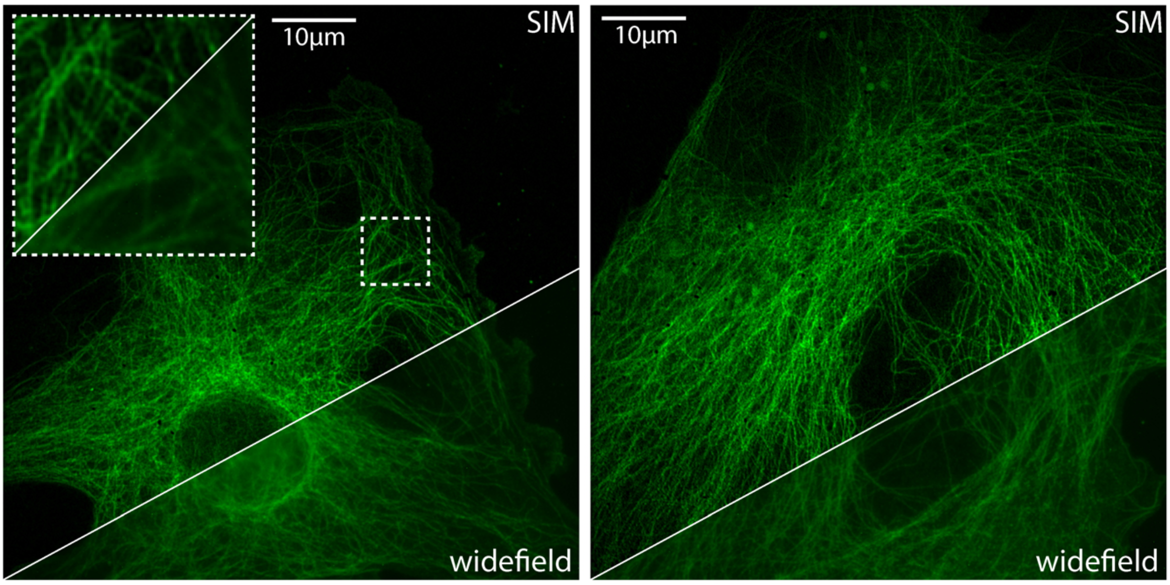


**Supplementary Figure 9: Wide-field and SIM images of microtubules with openSIM.**

Images of fixed bovine pulmonary artery endothelial cells, α-tubulin labeled with BODIPY. The inset represents a zoom in the area highlighted with a dotted square.
